# Bodily sensations in social scenarios: Where in the body?

**DOI:** 10.1101/441212

**Authors:** Giovanni Novembre, Marco Zanon, India Morrison, Elisabetta Ambron

**Author notes:** These authors contributed equally to this work (Giovanni Novembre, Marco Zanon).

## Abstract

Bodily states are fundamental to emotions′ emergence and are believed to constitute the first step in the chain of events that culminate in emotional awareness. Recent works have given fresh support to this view by showing that distinct topographical maps can be derived to describe how basic and more complex emotions are represented in the body. However, it is still unclear whether these bodily maps can also extend to emotions experienced specifically within social interactions and how these representations relate to basic emotions. To address this issue, we used the emBODY tool to obtain high-resolution bodily maps that represent the body activation and deactivation experienced by healthy participants in social scenarios depicting establishment or loss of social bonds. We show that clear patterns of activation/deactivation across the body emerge depending on the valence and on the characteristics of the single social scenarios, but also a common activation of head, chest and limbs across scenarios. Furthermore, we showed that maps related to complex social scenarios are strongly correlated with bodily states experienced in basic emotions, suggesting that the patterns of body activation/deactivation found for social scenarios might represent a combination of different basic emotions these experiences elicit. Our data tackle conscious emotional experiences and show for the first time how ′painful′ social events are felt on the body, providing findings that complement verbal reports and neuroimaging data on social rejection. Furthermore, they show that bodily feelings related to complex social scenarios, despite referring to unified independent emotional states, are also related to bodily correlates of basic emotions.

## INTRODUCTION

Humans continuously seek social affiliations. The establishment of new social bonds increases the chances of reproductive opportunities and more in general belonging to social groups results in undeniable advantages in terms of cooperation and survival strategies (1). Not surprisingly, the establishment of successful social connections has a positive impact on physical and mental health (2). On the contrary, social isolation is significantly associated with increased risks of mortality (3, 4), immune dysregulation (5), sleep fragmentation (6), and daytime fatigue (7). The effect of social bonding goes beyond the physical health of an individual, having important effects also on cognition and emotion regulation. For instance, social isolation is associated with drops in cognitive performances (8) and increased depression (9).

Everyday life experiences of social disconnection have been investigated in experimental settings by inducing transient feelings of social rejection, mostly using paradigms that were subsequently adapted to the MRI scanner (10). This line of research has revealed that even short experiences of social exclusion are perceived as extremely threatening, to the extent of being processed in the brain by parts of the same neural circuits involved in processing physical threats (11, 12). For instance, Novembre et al. (2015) showed that being excluded by other people in an online virtual game of catch a ball (Cyberball) activated in healthy volunteers the same brain regions that were recruited when painful electric stimulations where delivered to their own hands (rostral anterior cingulate cortex, posterior insular cortex and somatosensory cortex). In this regard, the similarity between the processing of social and physical threats points at the importance of social relationships for the survival of the individual, to the extent that they are processed as those stimuli attempting the physical well-being of the individual (13). Following this account, damages to social connections activate the same neural and physiological ′alarm system′ that responds to other relevant survival threats, such as the experience of physical harm (14). In support of this view, different studies have shown that experiences of social exclusion trigger a broad spectrum of physiological reactions, like drops in skin temperature (15), change in heart rates (16), and decreased interoceptive accuracy (17).

Taken together these findings suggest a close and direct association between affective processing of social bonds threats and body sensations. If on one hand this represents a novelty and a new area of investigation in social cognition, on the other hand the relationship between bodily states, feelings and emotions is a well-established topic and theoretically accounted by both classical (18) and more recent models of emotions (19, 20).

A study (21) has provided an important contribution to this debate, showing the existence of specific topographic representations on the body of primary emotions within the body. Specifically, Nummenmaa et al. demonstrated that maps of bodily sensations could be identified for different basic (e.g., fear and happiness) and complex (e.g., anxiety and love) emotions. Moreover, these maps were similar across different cultures and presentation modalities (i.e. video, words, vignettes), supporting the idea of a biologically-driven correspondence between visceral and somatic sensations and psychological emotional states (22).

Interestingly, a tight link between bodily sensations and emotional processing has been suggested also for experiences of social disconnection (13). Such experiences are usually defined as ′painful′, and felt emotions are often described in most of the languages by referring to physical terms such as ′broken heart′, ′hurt feelings′, or ′cutting stab′ (23). Within this set of literature, previous research has focused by most on investigating the overlap between social and physical pain in terms of neural substrates, while systematic investigation of bodily correlates of these emotions has been neglected. The present study aimed at filling this gap by investigating how sensations associated with social situations are represented in the body, in which pre-existing social connections are threatened or lost, or conversely when new bonds are built (13).

Specifically, we were interested in investigating whether participants represented body activation and deactivation for positive and negative social scenarios following specific patterns. To achieve this aim, we used the emBODY tool (Nummenmaa et al., (24) that allows to present online body silhouette and to obtain high-resolution bodily maps based on participants′ drawings on body silhouette. Participants were presented with eight matched negative and positive social scenarios (bereavement/birth, romantic rejection/acceptance, exclusion/inclusion, negative/positive evaluation) and they were asked to identify and highlight in the silhouette using a mouse the body activation and deactivation elicited by each scenario.

Second, we aimed at replicating Nummenmaa et al. (24) results with basic emotions (sadness, anger, fear, disgust, happiness, surprise) in out sample. In the light of the maps obtained by Nummenmaa et al., (2014a), we expected to find a clear dissociation between negative and positive social scenarios, with the former being mostly characterized by deactivation in the limbs (resembling maps of sadness and depression), and the latter mostly presenting activations spread throughout the body (recalling happiness and love maps).

Finally, a third aim of our work was to investigate the relationship between the body maps describing basic emotions and the one representing social scenarios. If the body representation of emotions elicited in social situations is related to the way we feel basic emotions, we expected positive correlations between bodily representation of basic emotions and social scenarios only of similar emotional valence. Specifically, we expected that maps depicting negative social scenarios would show positive correlations with negative basic emotions (i.e. sadness, anger, fear, disgust), whereas maps of positive social scenarios would be positively correlated with positive basic emotions (i.e., happiness).

## METHODS

### Participants and experimental procedure

The present study was carried out as web survey and promoted through discussion on online forums and social networking. The website registered 247 attempts to undertake the experiment but only 95 participants underwent the whole experimental session. The web survey was composed of three parts: (1) collection of demographic information; (2) experimental tasks (body localization of social scenarios first and basic emotions after); (3) presentation of a series of questionnaires. These were: (1) Beck depression inventory (BDI; (25–27)), a self-reported questionnaire with 13 items in which participants were asked to indicate a sentence among 4 which best described their actual emotional state; (2) Bermond-Vorst Alexithymia Questionnaire version B, (BVAQ-B; Vorst and Bermond 2001 (28), a 20-item-scale measuring emotional awareness, considered as a proxy for alexithymia; (3) A 8 item post-task questionnaire (PTQ) in which participants used a visual analog scale ranging from 0 to 10 to rate the intensity of the social scenarios presented during the task.

One participant who did not provide demographic information and three participants who declared to be under psychiatric treatment were excluded from the final sample. Therefore, the final sample was composed of 91 Italian participants (65 females, age *M*= 30.1, *SD*= 9.1). None of the participants reported severe depressive symptoms. In order to estimate a minimum sample size for obtaining meaningful maps for each emotion, we used G*Power software (29) to run an *a priori* power analysis with the following inputs: 2-tailed one sample t-test as statistical test, effect size of 0.5, alpha level of 0.05, and power of 0.95. This resulted in an estimation of at least 54 participants for the present study.

### Experimental tasks

#### Body localization task for social scenarios

The emBODY tool (21) was used also to localize bodily sensations in social scenarios. In each trial, participants viewed a short text presented in the center of the screen, with two silhouettes placed on the right and left sides (Fig 1). The sentence described either a negative or positive social scenario such as: bereavement, romantic rejection, social exclusion, and negative evaluation (negative social scenarios); birth of a son, romantic acceptance, social inclusion, and positive evaluation (positive social scenarios, see Table 1). For each scenario, participants were asked draw on both silhouettes the portions of the body where they felt activation (left silhouette) and deactivation (right silhouette) when facing the event described in the sentence.

**Fig 1.**
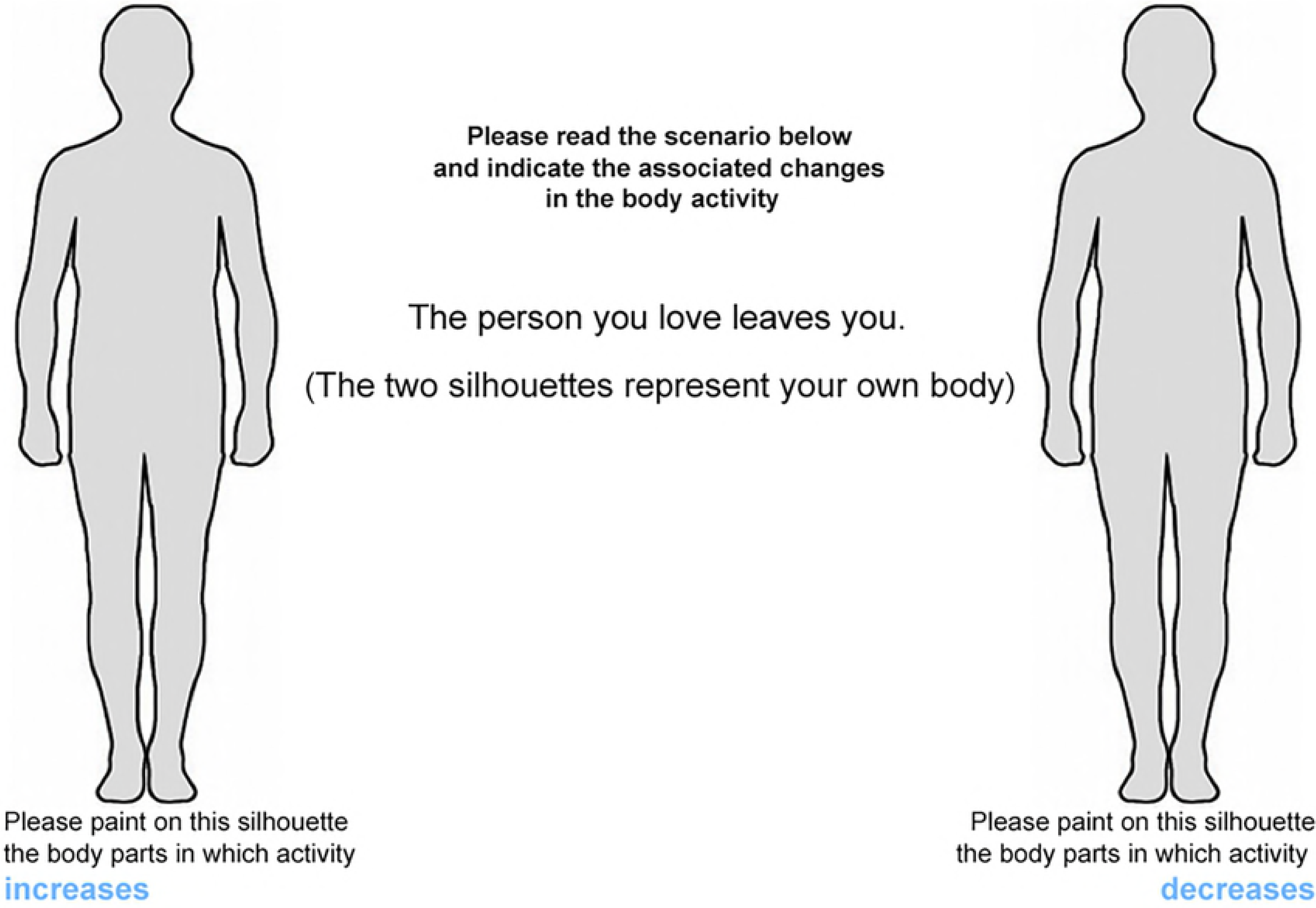
Representation of a single trial (romantic rejection, self-condition) as seen from the participant′s perspective. The specific task instructions are shown centrally on top, whereas the description of the scenario is shown in the center, between the two silhouettes.

**Table 1.** List of social scenarios presented to participants in the social scenario localization task.

|  |
| --- |
| **Bereavement/Birth** |
| The death of your mother |
| The birth of your child |
| **Romantic rejection/acceptance** |
| The person you love breaks up with you |
| The person you like tells you (s) he is in love with you |
| **Exclusion/Inclusion** |
| You are not invited to a dinner to which you expected to be invited |
| You are invited to a party to which you did not expect to be invited |
| **Negative/Positive evaluation** |
| Your friends disapprove your way of being |
| Your friends praise you for your personality |

In each scenario, the participant was the subject of the event in the sentence (e.g., *′the person you love breaks up with you′*) and was asked to focus on his/her own sensations when imagining being in that situation. Each sentence was presented once in a random order.

#### Body localization task for basic emotions

We used the same basic task as Nummenmaa et al.′s study (24). Participants were instructed to localize the subjective bodily sensations evoked by positive and negative basic emotions such as sadness, anger, fear, disgust, happiness, and surprise. Emotions were presented as single words in the center of the screen and participants were asked to paint the parts of the body where they feel usually an increase or decrease of activation when experiencing the depicted emotion. This task consisted of 6 trials presented in a pseudorandom order.

### Data pre-processing and analysis

Data preprocessing and analysis was carried out using custom scripts on the MATLAB platform (R2010a, The Mathworks, Inc., USA).

#### Localization maps

First, we explored the topographic representation of each basic emotion and social scenario across participants, using the Nummenmaa and co-workers′ methodology (2014) to reconstruct bodily sensation maps (BSMs) from data collected during the web survey. For each participant, a single BSM comprising 50364 pixels was obtained for each basic emotion and social scenario, with activation and deactivation coded respectively as positive and negative values, and subject-wise normalized to the maximum color intensity provided by the participant, such that final paint intensity ranged from 0 to 100. For each basic emotion and social scenario, statistically significant activations and deactivations were assessed by means of mass univariate *t*-tests: a one-sample *t*-test against zero was performed for each pixel within a BSM, resulting in a statistical *t*-map. To account for multiple comparisons, each statistical map was then thresholded using the False Discovery Rate (FDR) correction (significance level = 0.05).

To further investigate the patterns of bodily sensations, we divided the body silhouette in 5 parts, corresponding to the head, the chest, the abdomen, the upper limbs (arms and hands), and the lower limbs (legs and feet) and averaged the participant’s color intensity values in each of these parts. Significant activations and deactivations in each body part were assessed with a one-sample *t*-test against zero with the initial significance level (*p* = 0.05) Bonferroni-corrected for multiple comparisons.

#### Overlap maps

We tested for commonalities across BSMs using masks made of the significant pixels obtained in the one-sample *t*-tests analysis. Specifically, we created binary masks for each basic emotion and social scenario converting significant and not significant pixels in values of 1 and 0, respectively. Masks were then overlapped and the resulting map was color-coded such that each color corresponded to the number of maps that showed a significant value for that pixel. Maps of overlap were created for positive and negative social scenarios referred to the self or to the other person.

#### Correlation matrices

In order to investigate the relationship between body representations of social scenarios and basic emotions, we correlated the mean intensity of each body part (i.e., head, chest, abdomen, arms and legs) between social scenarios and basic emotions and among basic emotions. Significant level for all correlations was set at *p* = 0.05 and the FDR correction was applied.

## RESULTS

### Body localization maps

#### Social scenarios

For positive social scenarios, participants mainly identified body activations, in particular in the chest, abdomen and head regions, whereas for negative scenarios participants reported both activations and deactivations, with the former localized primarily in the head and chest regions, and the latter related to the legs and arms (Fig 2).

**Fig 2.**
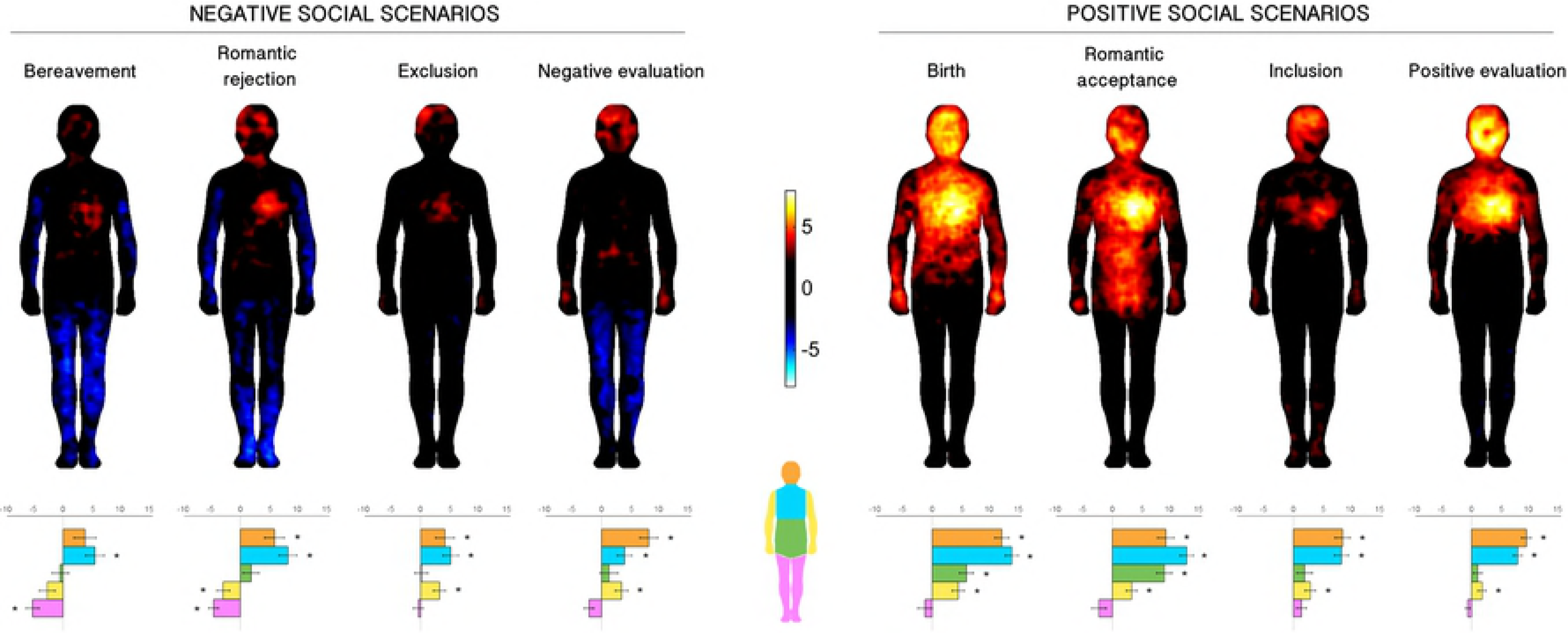
BSMs associated with social scenarios. Left panel: negative scenarios. Right panel: positive scenarios. Regions of increased and decreased activation are depicted together on the same maps, respectively in warm (0 to 8) and cool colors (0 to − 8). Maps are thresholded at p < 0.05, FDR-corrected. The bar plots represent the intensity for each body part (orange = head, blue = chest, green = abdomen, yellow = arms, violet = legs). Asterisks indicate values that differ from zero (*p* < 0.05, Bonferroni-corrected).

One sample t-tests confirmed these observations (see bar graphs below each BSMs in Fig 2) as all positive scenarios presented significant (p < 0.05, Bonferroni-corrected) positive values in the head, chest, abdomen, and arms, suggesting a significant activation of these areas. The activation in the abdominal region characterized birth and romantic acceptance, while inclusion and positive evaluation did not present any significant activation in this region.

Furthermore, statistical tests confirmed that differently from positive scenarios, negative scenarios were characterized by mixed patterns of activations and deactivations. For this latter set of scenarios, participants reported activations in the head, chest and arms, and deactivations in the limbs. Specifically, bereavement was characterized by activation of the chest and deactivation of the arms and legs. Romantic rejection presented a similar pattern, with significant activation of the head and chest. As for exclusion, participants identified activation in the head, chest and arms (hands). Finally, a significant activation in head and chest was observed for negative evaluation, along with an increase of activation in the arms (hands).

Since differences in the ability to identify and describe their own affective states may influence participants′ performance in the task, we also investigated the difference across maps between three subgroups of participants defined according to their scores in the BVAQ-B questionnaire (non-alexythymic n=39, borderline n=22, alexithymic n=30). We first obtained BSMs for each emotion independently for each group with the same approach explained before for the whole sample, and then we ran a one-way ANOVA to explore the main effect of the group. This analysis did not show any significant difference between the subgroups.

### Overlap maps

Most of the maps of the positive social scenarios overlapped in the face and chest areas and were mainly defined by activation. For negative social scenarios, most of the activations across the maps overlapped in the face, head and chest area around the heart region, while deactivation overlapped in upper and lower limbs in most of the maps (Fig 3).

**Fig 3.**
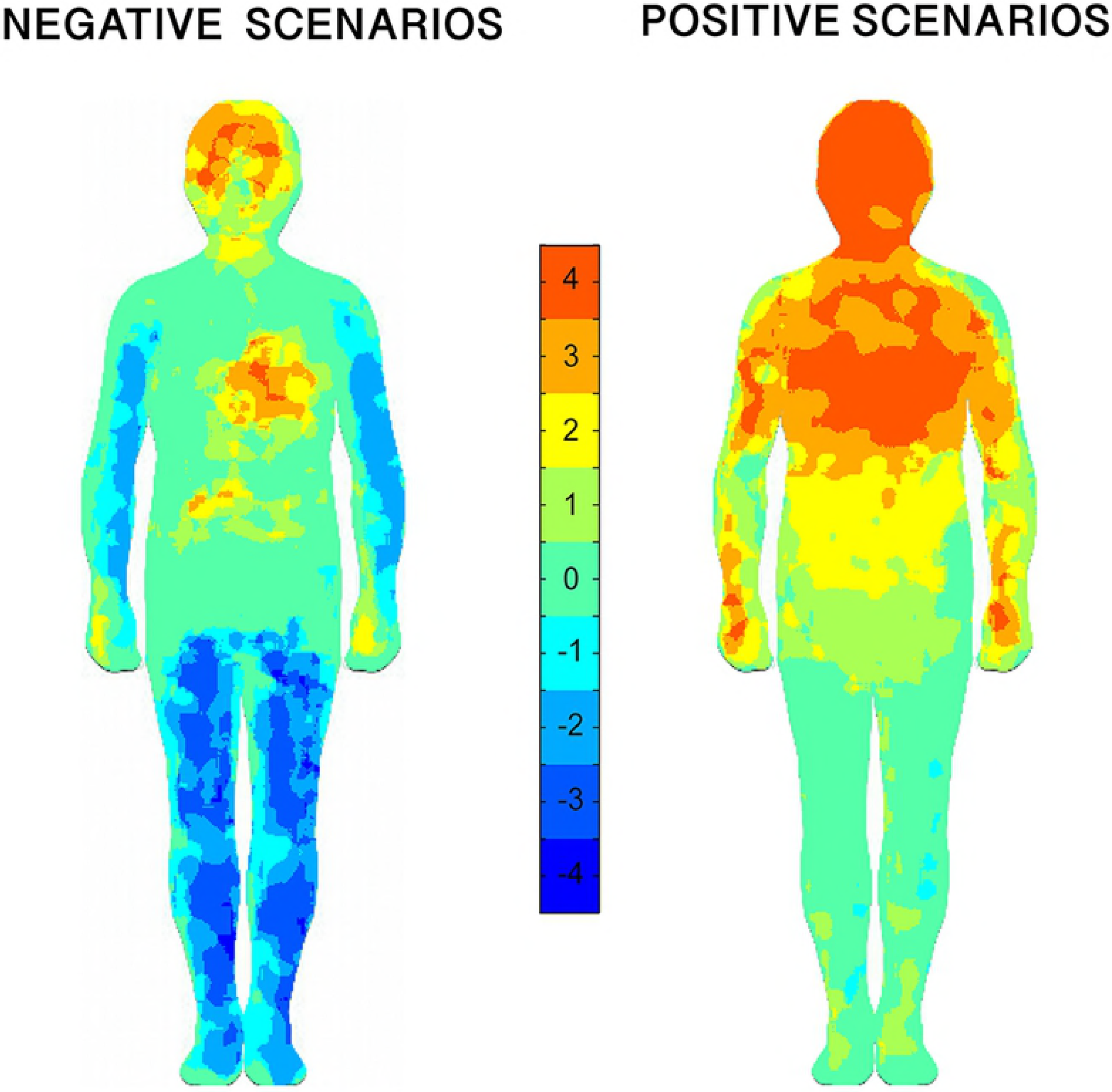
Overlaps of masks computed from negative and positive scenarios. Masks included only significantly activated and deactivated pixels. The colormap indicates the number of overlapping significant pixels.

#### Basic emotions

We replicated the results of Nummenmaa et al. (2014). Indeed, basic emotions evoked specific patterns of activations and deactivations in the different body parts (see bar graphs below BSMs in Fig 4). One-sample *t*-tests against zero showed that sadness was the only emotion that presented significant (p< 0.05, Bonferroni-corrected) deactivations, specifically in the arms and legs. Most of the other emotions showed activations in the head (except for fear) and the chest (except for disgust). Anger was further characterized by activations of the arms, particularly in the hands, whereas happiness presented diffuse activation of the chest that crossed in the arms and the upper abdomen.

**Fig 4.**
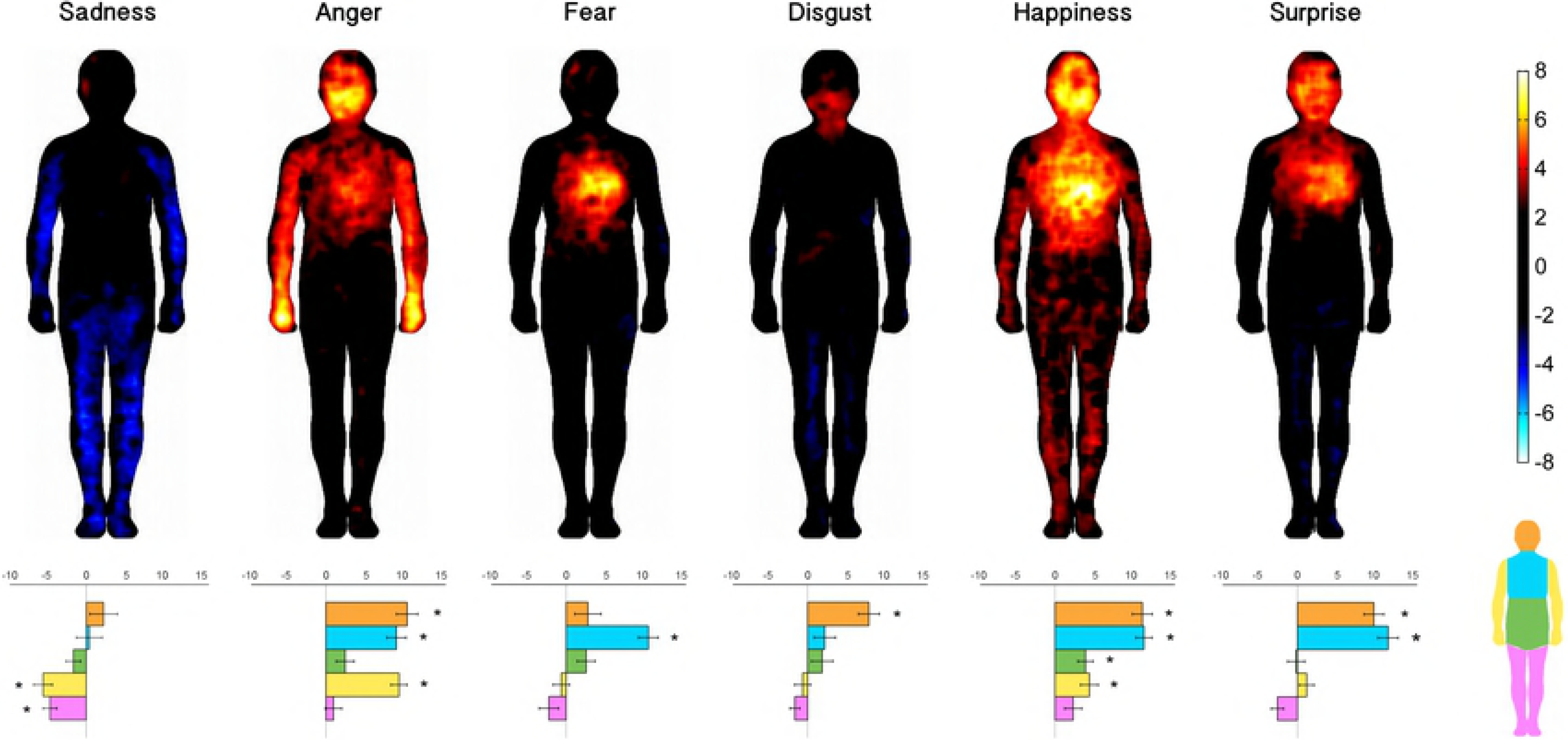
BSMs associated with basic emotions. Regions of increased and decreased activity are depicted together from warm (0 to 8) to cool colors (0 to − 8). Maps are thresholded at p < 0.05, FDR-corrected. The bar plots represent the intensity the for each body part (orange = head, blue = chest; green = abdomen, yellow = arms, violet = legs). Black asterisks indicate values that differ from zero (p < 0.05, Bonferroni-corrected).

#### Correlation matrices

Here we report the correlation indexes between body-part mean intensities reported during imagined experiences of each social emotion and the same values associated with all basic emotions (Fig 5). All social scenarios showed only positive correlations with basic emotions. The four negative social emotions (left column) correlated mostly with negative nonsocial emotions (left part of the matrices: sadness, anger, fear, disgust), whereas the four positive social emotions (right column) correlated highly with neutral/positive basic emotions (right part of the matrices: happiness, surprise).

**Fig 5.**
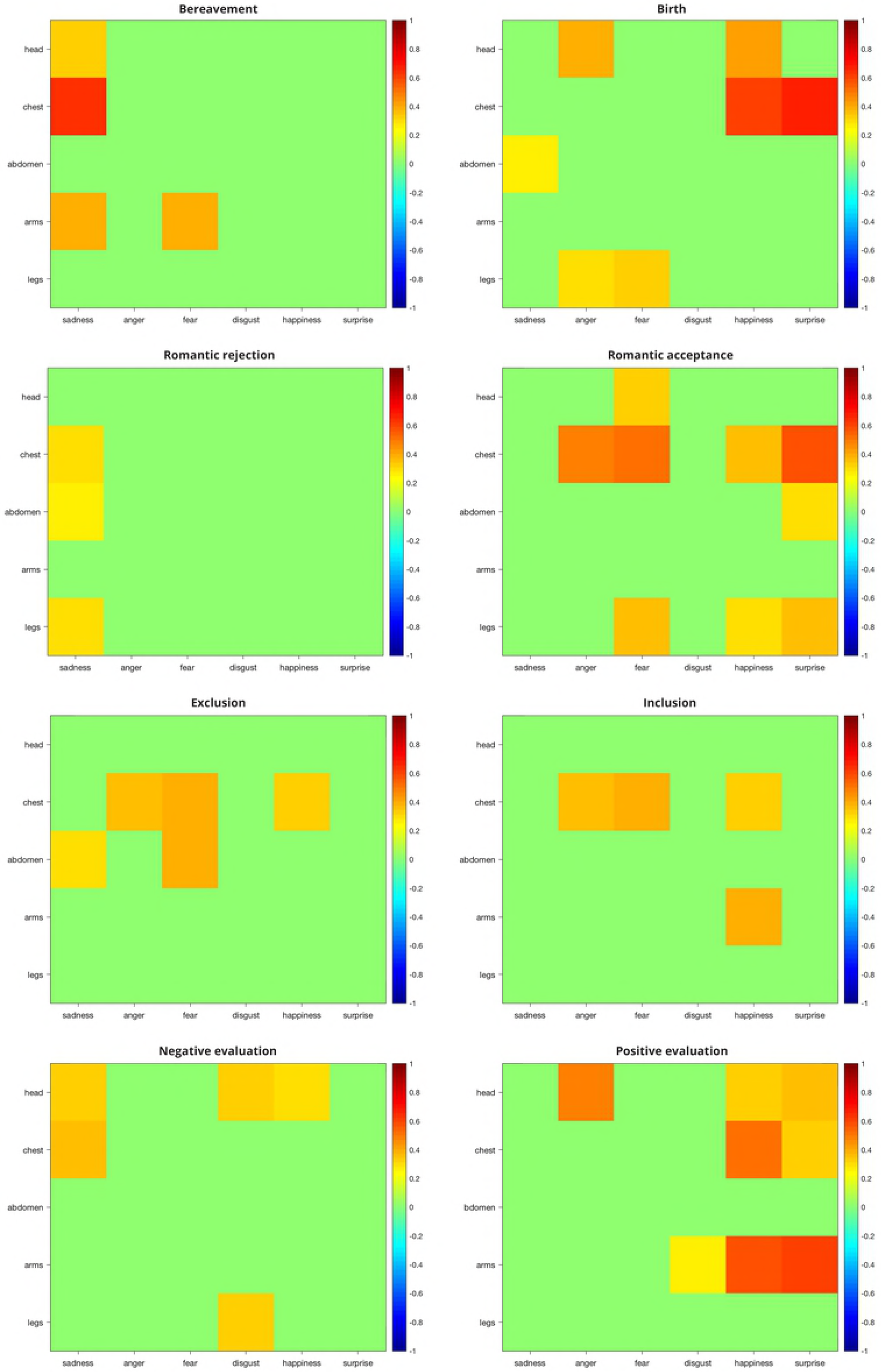
Correlational matrices between social and basic emotions. Each panel depicts the correlation matrix of a social emotion with the six basic emotions (x-axis), separately for each body part (y-axis). Positive and negative Spearman′s coefficients are indicated respectively in warm and cool colors. Only significant coefficients are reported (p < 0.05, FDR-corrected).

Looking at negative emotions, bereavement correlated mainly with sadness in head, chest, and arms; romantic rejection correlated with sadness in chest, abdomen and legs; social exclusion presented correlations with sadness, anger and fear in the torso; finally, negative social evaluation correlated with sadness in the head and chest, and disgust in head and legs.

Similarly, positive social scenarios showed subtle differences within a consistent general pattern. In particular, birth correlated with positive basic emotions especially in the head and chest; romantic acceptance showed significant correlations with positive basic emotions in chest, abdomen, and legs; inclusion correlated with happiness in the chest and in the arms; finally, positive evaluation showed significant correlations with positive basic emotions in the whole body in the head, chest and arms.

Interestingly, although negative and positive social scenarios correlated by most with basic emotions of the same valence, they also presented significant positive correlations with basic emotions of the opposite valence. In particular, negative social scenarios correlated with happiness (in romantic rejection and negative evaluation), whereas all positive social scenarios correlated with anger, and most of them with sadness and fear. Positive evaluation also correlated with disgust in the arms.

Therefore, in order to disentangle meaningful and spurious correlations, we also calculated the same correlation coefficients among basic emotions (Fig 6). Basic emotions correlate not only with other basic emotions of the same valence, but also with emotions of opposite valence, suggesting a similar activation of body parts across the whole spectrum of emotional valence. For instance, happiness and surprise were positively correlated with anger and fear in the chest.

**Fig 6.**
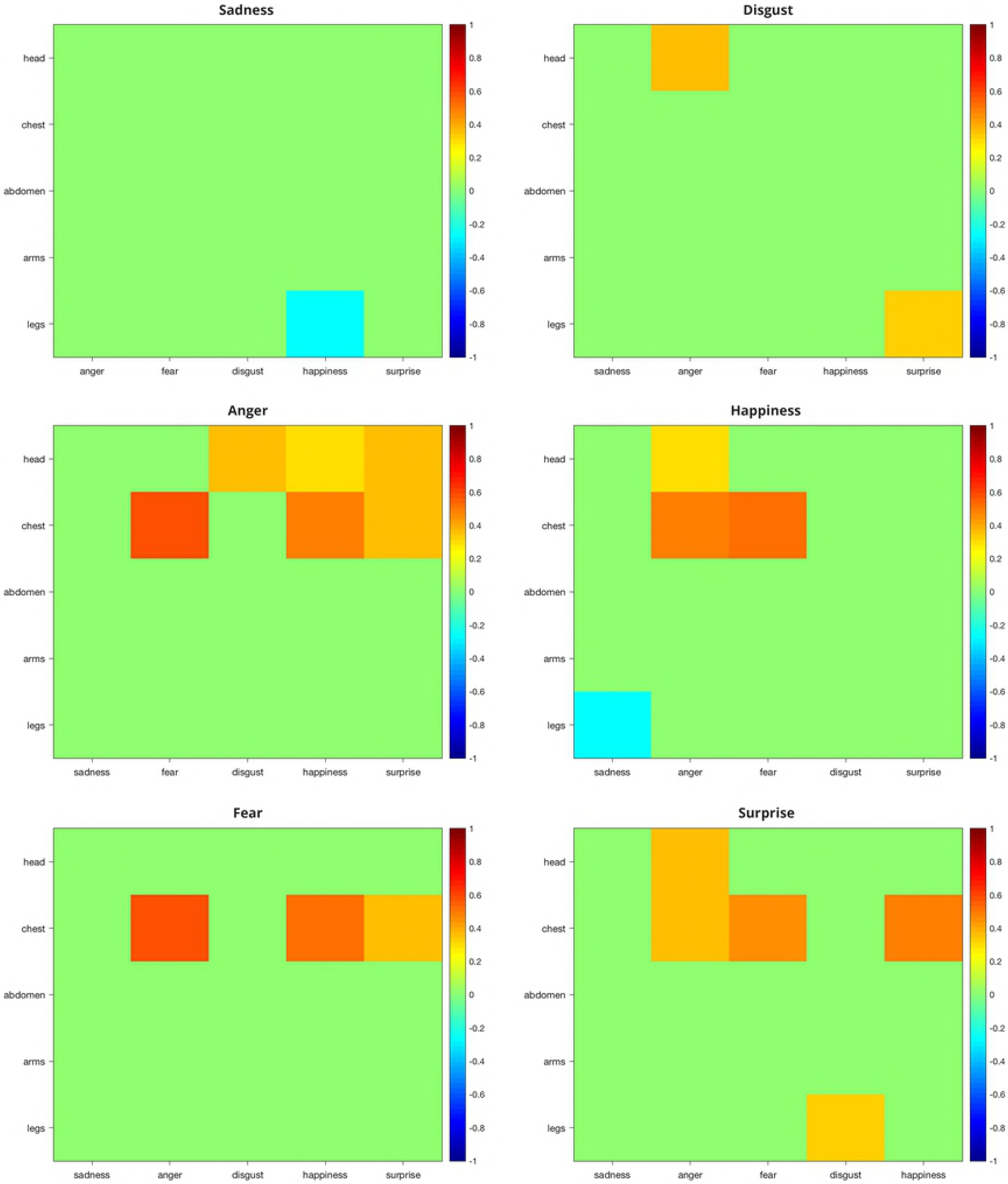
Correlational matrices between pairs of basic emotions. Each panel depicts the correlation matrix of a basic emotion with all the other basic emotions (x-axis), separately for each body part (y-axis). Positive and negative Spearman′s coefficients are indicated respectively in warm and cool colors. Only significant coefficients are reported (p < 0.05, FDR-corrected).

#### Post-task questionnaire

Differences between the ratings of the intensity of social scenarios were tested with a repeated-measures ANOVA with *scene* (4 scenarios) and *valence* (positive and negative) as within-subject factors. Results showed a main effect of *scene* (*F*_3,88_ = 102.836, *p* < 0.001) and a significant interaction between *scene and valence* (*F*_3,88_ = 4.995, *p* = 0.003). Participants rated as more intense the combination of birth/bereavement and of romantic acceptance/rejection scenarios than both positive/negative evaluation and inclusion/exclusion scenarios (*p* < 0.001, Bonferroni-corrected for all the comparisons) (see Table 2).

**Table 2.**
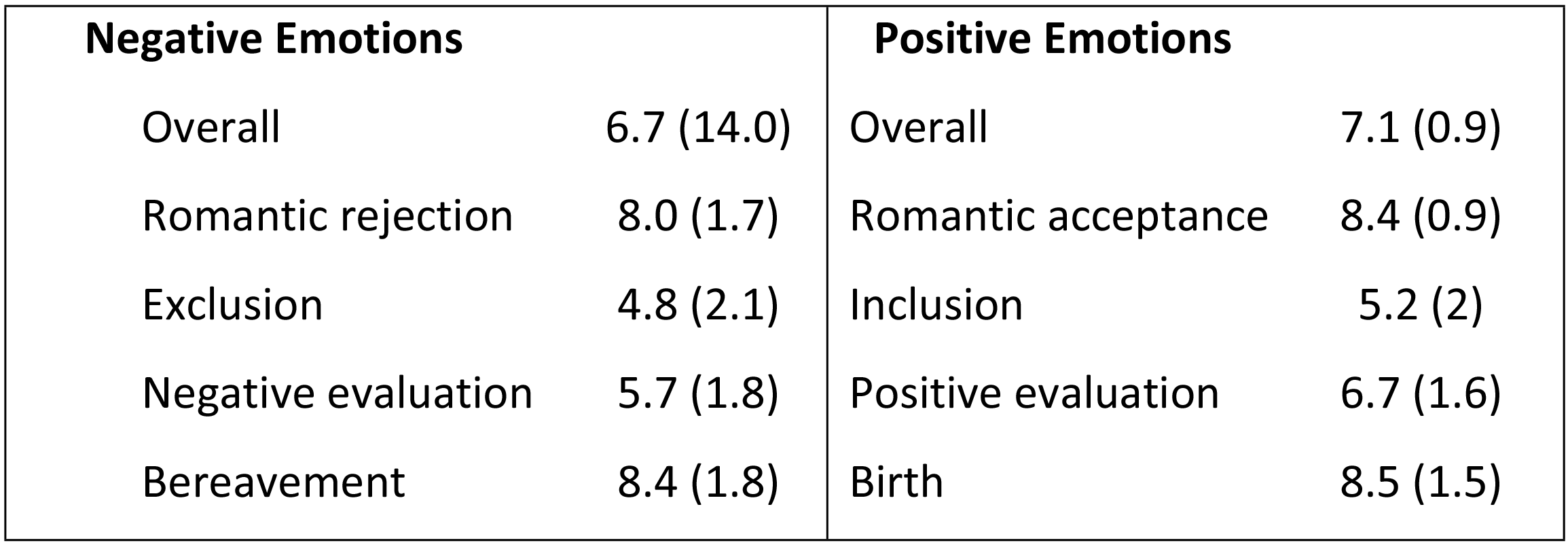
Mean (SD) explicit ratings of emotional scenarios.

| Negative Emotions |  | Positive Emotions |  |
| --- | --- | --- | --- |
| Overall | 6.7 (14.0) | Overall | 7.1 (0.9) |
| Romantic rejection | 8.0 (1.7) | Romantic acceptance | 8.4 (0.9) |
| Exclusion | 4.8 (2.1) | Inclusion | 5.2 (2) |
| Negative evaluation | 5.7 (1.8) | Positive evaluation | 6.7 (1.6) |
| Bereavement | 8.4 (1.8) | Birth | 8.5 (1.5) |

### DISCUSSION

Social bonds are crucial for human lives such that the foundation or disruption of social connections are associated with emotional experiences that are described in terms of perceived changes in the body (23). In our study, we detailed the representations of the perceived bodily changes elicited by social scenarios in which social bonds were created (e.g., romantic acceptance) or broken (e.g., romantic rejection).

By using the emBODY tool (Nummenmaa, 2014), we focused on the bodily sensations associated with a specific subset of emotional experiences, namely social scenarios, that involve the constitution or dissolution of social bonds, with two main goals: (1) to provide a qualitative description and topographic representation of these social scenarios and test their communalities; (2) to replicate previous findings on basic emotions (sadness, anger, fear, disgust, happiness, surprise) in our sample; (3) to investigate the relationship between the body maps describing basic emotions and the ones representing social scenarios.

First, we showed that social scenarios are associated with changes in activation and deactivation of specific body regions likewise for basic emotions (24), so that topographic representations can be derived for specific social scenarios (bereavement/birth, romantic rejection/acceptance, exclusion/inclusion, negative/positive evaluation). Furthermore, we showed that these topographic maps present more commonalities than differences. Communalities across maps were observed when looking at single BSMs maps (Fig 2), but even more striking when looking at the overlapped maps (Fig 3). In particular, activation of the chest and face areas and deactivation of the limbs represented the most consistent pattern across all emotions. In addition, bodily activation-deactivation patterns were also modulated on the basis of the valence of the emotions. Indeed, positive emotions were felt by most as activation of the head/face and the chest. This pattern was also observed with negative emotions but with less intensity and extension, so that activations were restricted to the upper part of the head and to the heart region. At the same time, negative emotions were represented with the deactivation of the limbs, especially of the legs. These valence-specific patterns were confirmed when computing and analyzing the mean intensity values for the 5 body parts (the head, the chest, the abdomen, the upper limbs, and the lower limbs).

The consistent activation observed in the head area suggests that people associate social scenarios with high-level mental processing. This is not surprising as the social scenarios depicted situations of creation or disruption of social connections that require high level of social cognitive processing, as evaluation of personal states in relation to others (e.g., the break-up with the partner) (30). Notably, the patterns of activation and deactivation observed for the head region are the most heterogeneous and rich across the different body areas, as they reflects changes not only in the brain, but also in the face with all its parts. Some scenarios enhanced the identification of very specific areas within the face areas. For example, for negative scenarios like bereavement and romantic rejection participants painted well-defined blobs around the eyes, possibly indicating lacrimation, whereas in all positive scenarios clear activation blobs cover the cheeks areas, reflecting probably blushing and facial muscular activation.

In the same fashion, the persistent activation reported in the heart/chest area across positive and negative scenarios is not surprising as it might reflect the link between the representation of these scenarios and the increase in heart rate, commonly experience for both positive and negative emotional states. Accordingly, previous work has shown the association between increased heart rate and both positive and negative emotions like anger, anxiety, fear and sadness on one hand, and with happiness, joy and anticipated pleasure (31). Also, increased heart rate has been found as one of the main response of distress following social rejection (16) and in social situations more in general (32–34).

While a common factor across maps is the representation of an activation of the head and the heart region, a more connotative distinction between positive and negative scenarios can be identified when looking at the upper and lower limbs. As shown in the overlap maps, deactivation emerges as a key feature of negative scenarios and localized in both upper and lower limbs, whereas activation in upper limbs distinguishes positive scenarios. These distinctive body-representations seem to reflect physiological and sensorimotor changes related to emotional experiences of opposite valences and could represent approach and avoidance tendencies (35). We could speculate that the higher activity in the arms for positive scenarios may reflect approach-oriented actions, for instance to lift the newborn up in the birth scenario or to hug the partner in the romantic acceptance. On the other hand, negative emotions may be rather associated with immobility and avoidance, resembling the deactivation found in the depression map obtained by Nummenmaa and colleagues (2014).

Emotional experiences in general are strongly associated with bodily sensations reflecting changes in neuroendocrine, skeletomuscular, and autonomic nervous systems in response to internal or external stimuli (20, 36). Rather than being generic, the bodily changes associated with emotional states have distinct topographical signatures that are mapped in the body. In fact, Nummenmaa et al. (2014) showed that basic and nonbasic emotions are associated with distinctive, yet partially overlapping representations of the body activation and deactivation (21). In the present study, we replicated the results for the basic emotions, obtaining maps that are comparable to the ones presented in Nummenmaa′s study. For instance, we found that emotions like anger and happiness were associated with greater activity in the upper limbs, whereas decreased limb activity characterized sadness. Furthermore, disgust was associated with higher activation represented in the throat region, whereas anger, fear, happiness and surprised were confirmed to be the emotions presenting the higher activity in the upper chest, probably reflecting increased breathing and heart rate.

A final aim of this study was to investigate whether bodily representation of social scenarios could result from the simultaneous contribution of basic emotions. The correlation analyses between the intensity of the basic emotion and social scenarios in each body part showed a strong association between emotions of a similar valence. Negative social scenarios, all having in common the experience of a threat to or an actual loss of a social bond, correlated with negative nonsocial emotions like sadness, anger, fear, and disgust. Feelings of social exclusion have been reported in terms of anger, anxiety, depression and shame (37, 38). This association between social scenarios and basic emotions is coherent with classical theories positing that complex emotions emerges from primary emotions in the course of socialization. As suggested for guilt, shame and pride, that are thought to gain their emotional tone by virtue of their linkage with fear, anger and happiness respectively, complex emotions regarding social (dis-) connection scenarios can attain their status of emotions through coupling with basic emotions (39, 40). Nonetheless, given the correlational nature of these data, further studies should be devoted to explore the possible causal relations and systematically investigate whether this association is not confined to a representational level only but if these emotions enhance also similar autonomic responses.

The present study provides a starting point to better understand how emotions in social scenarios are represented in the body, but it presents some limitations. First, our work is based upon self-reported data regarding where activation and deactivation related to these scenarios are represented in the body, since our main interest was how people represent social-scenarios related emotions in the body, so we did not directly investigate the actual change in physiological state. Thus, future research should investigate more systematically whether the way people perceived bodily changes related to social situations reflect actual physiological changes (i.e. changes in heart rate, blood pressure, body temperature, respiration rate, etc.). Second, we did not test for the possible effects of the interoceptive ability (41). Indeed, variation across individuals in the ability to accurately detect bodily changes may have influenced the performance in our task and future studies should address this issue.

## CONCLUSIONS

In the present work, we used a high-resolution and language-independent tool (emBODY) to obtain bodily maps of emotions linked to establishment and unwanted disruption of social bonds. Results show that clear patterns of activation/deactivation across the body emerge depending on the valence and on the characteristics of the single scenarios, but they assign a central role to psychological and physiological processes associated with head, chest and limbs.

We think that these maps can be of great help to social rejection researchers, in that they bridge the gap between the psychological and physical components of socially painful experiences.

## Acknowledgements

We would like to thank Enrico Glerean providing us with the emBODY tool and for his support in the use of the program. We would also like to thank Alessio Isaja for his technical support.

